# The mechanisms of siRNA selection by plant Argonaute proteins triggering DNA methylation

**DOI:** 10.1101/2022.06.01.494257

**Authors:** Wei Liu, Keisuke Shoji, Masahiro Naganuma, Yukihide Tomari, Hiro-oki Iwakawa

## Abstract

The model plant *Arabidopsis thaliana* encodes as many as ten Argonaute proteins (AGO1–10) with different functions. Each AGO selectively loads a set of small RNAs by recognizing their length and 5′ nucleotide identity to properly regulate target genes. Previous studies showed that AGO4 and AGO6, key factors in DNA methylation, incorporate 24-nt small-interfering RNAs with 5′ adenine (24A siRNAs). However, it has been unclear how these AGOs specifically load 24A siRNAs. Here, we biochemically investigated the siRNA preference of AGO4, AGO6 and their chimeric mutants. We found that AGO4 and AGO6 use distinct mechanisms to preferentially load 24A siRNAs. Moreover, we showed that the 5′ A specificity of AGO4 and AGO6 is not determined by the previously known nucleotide specificity loop in the MID domain but rather by the coordination of the MID and PIWI domains. These findings advance our mechanistic understanding of how small RNAs are accurately sorted into different AGO proteins in plants.

## INTRODUCTION

Eukaryotes produce various types of small RNAs about 20–30 nucleotides in length, such as microRNAs (miRNAs) and small interfering RNAs (siRNAs), that function in diverse biological processes including regulation of gene expression and defense against viruses and transposable elements (TEs). For these functions, small RNAs need to form an effector complex named RNA-induced silencing complex (RISC) with an Argonaute (AGO) family protein. RISC assembly consists of two steps: small RNA loading and RISC maturation (1). In general, small RNAs are first loaded into AGO in a duplex form, consisting of a guide strand and a passenger strand, with the help of the HSP70/90 chaperone machinery (2–4). After loading of small RNA duplexes, the passenger strand is ejected from AGO, while the guide strand remains in AGO to form mature RISC. Subsequently, RISC binds target mRNAs via base complementarity to the guide strand and silences them by endonucleolytic cleavage, mRNA decay, or translational/transcriptional repression (1).

AGO consists of four domains (N, PAZ, MID, and PIWI) and two linkers (L1 and L2). The catalytic activity of RISC relies on the PIWI domain, which forms an RNase H-like fold (5–8). The PIWI domain and the MID domain form a pocket that recognizes the 5′ end of the guide strand. The MID domain possesses the nucleotide specificity loop, which is thought to discriminate the 5′ end nucleotide of the guide strand (9).

Many organisms possess multiple AGOs with different functions. To achieve their diverse regulatory functions, small RNAs must be sorted into the correct AGO proteins. Small RNAs with a different nucleotide at the 5′ end are sorted into different AGOs in plants (10–12). The model plant *Arabidopsis thaliana* encodes as many as ten AGOs (AGO1–10). AGO1, AGO2, and AGO5 preferentially load 21-nt small RNAs bearing a uridine (U), adenine (A), and cytosine (C) nucleotide at the 5′ end, respectively. Previous structural studies and binding analyses showed that the nucleotide specificity loops in the MID domain of these plant AGOs recognize the specific nucleoside monophosphates (13), suggesting that the nucleotide specificity loop is responsible for small RNA sorting in plants.

Plant 24-nt siRNAs induce RNA-directed DNA methylation (RdDM) by associating with AGO4 clade proteins, including AGO4, AGO6 and AGO9, which are differentially expressed in a tissue-specific or locus specific manner (14–20). The 24-nt siRNAs are processed by DICER-LIKE3 (DCL3) from ∼38-nt double-stranded RNA (dsRNA) precursors, named P4RNAs or P4R2 RNAs, which are synthesized by DNA-dependent RNA polymerase IV (Pol IV) and RNA-dependent RNA polymerase 2 (RDR2) (Figure 1A) (19). Sequencing of total small RNAs in plant cells showed that about 60% of 24-nt siRNAs have a 5′ A (18). This enrichment of 24A siRNAs can be explained by the biased selection of transcription start site (TSS) by Pol IV and the substrate specificity of DCL3: Pol IV TSSs exhibit preference for A or G at the +1 position (21), and DCL3 prefers to cleave dsRNA precursors from the end with 5′ A and/or unstable terminus (22–25). On the other hand, sequencing of small RNAs loaded into AGO4/6 showed a slightly higher percentage of 24A siRNAs than total small RNAs (18), suggesting that 24A siRNAs are further enriched in the loading step. However, to what extent AGO4 and AGO6 actively select 24A siRNAs during the loading step remains to be determined.

**Figure 1.**
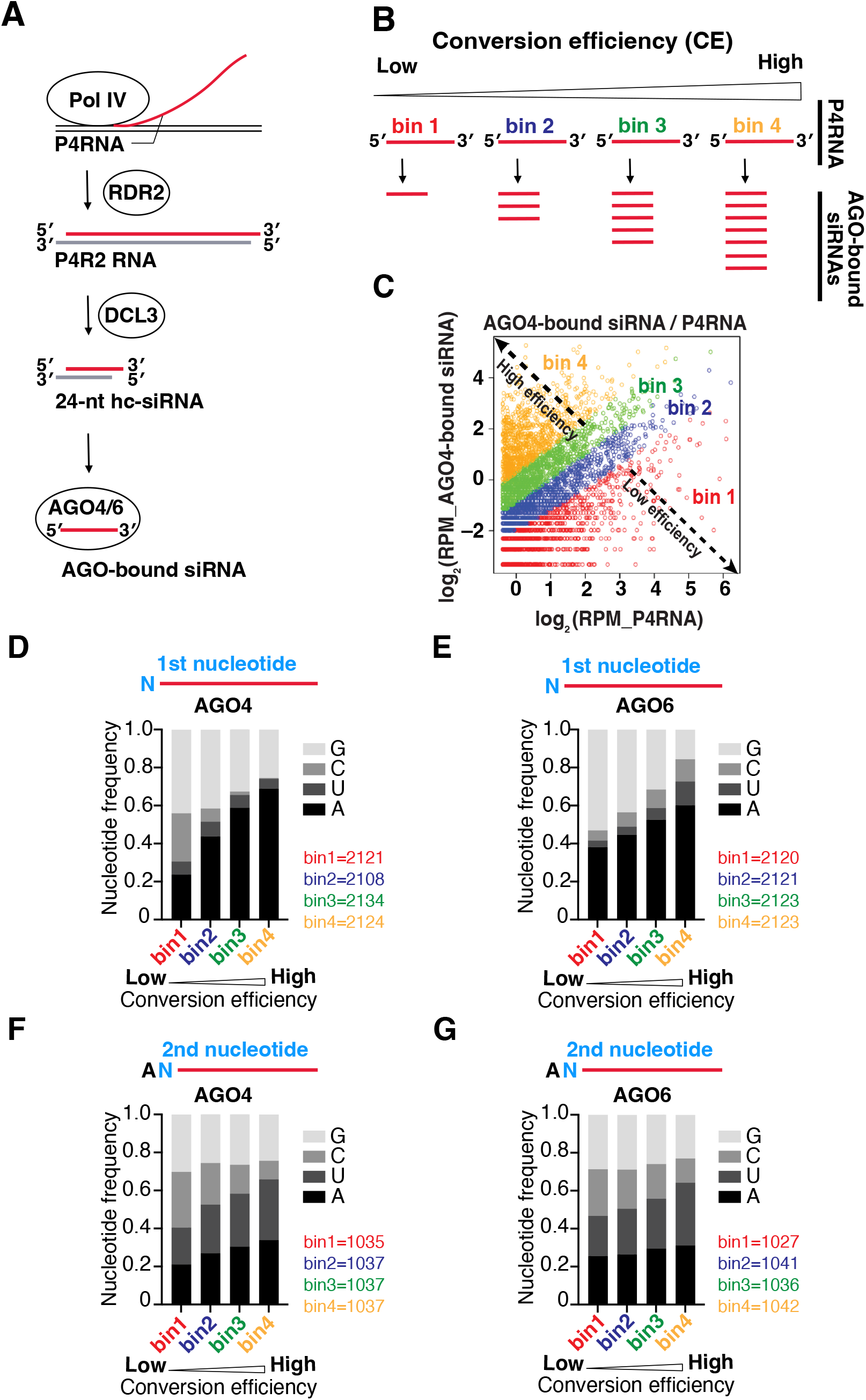
P4RNAs with 5′ A and/or unstable ends are successfully converted into mature siRNAs. (A) Schematic of 24-nt heterochromatic-siRNA biogenesis in *Arabidopsis*. P4RNA, a 30–40 nt single stranded RNA is first transcribed by Pol IV. Subsequently, a complementary strand is synthesized by RDR2, forming a long double-stranded RNA (dsRNA) with an extra nucleotide at the 3′ end of RDR2 strand. The dsRNA is subsequently diced by DCL3 into a 24-nt siRNA duplex, which is loaded into AGO4 or AGO6 to form RISC. The passenger strand is cleaved and ejected by AGO, while the guide strand remains in mature RISC. (B) P4RNAs are classified into four bins according to conversion efficiency (CE), from lowest bin 1 to highest bin 4. P4RNAs with higher CE produce more siRNAs. (C) An example scatter plot of AGO4-bound siRNAs and P4RNAs. Each spot represents one P4RNA. All P4RNAs were grouped into four bins with CEs ranging from low (bin 1) to high (bin 4). (D and E) First nucleotide frequency of AGO4- (D) and AGO6- (E) bound small RNAs. Number of P4RNAs in each bin is shown on the right side. P4RNAs with higher CE tend to have a stronger 5′ A bias in both AGO4 and AGO6. (F and G) Second nucleotide frequency of AGO4- (F) and AGO6- (G) bound 5′ A small RNAs. Number of P4RNAs in each bin is shown on the right side. P4RNAs with higher CE tend to have an A/U nucleotide compared to a C/G nucleotide at the g2 position in both AGO4 and AGO6.

Here, using an *in vitro* RISC assembly system and bioinformatics approaches, we quantitatively investigated the siRNA preference of AGO4 and AGO6. We showed that both AGO4 and AGO6 preferentially load 24A siRNAs but do so by distinct mechanisms. AGO4 selects 24A siRNAs via a combination of a moderate 5′ purine preference and a preference for unstable base-pairing at the 5′ end, whereas AGO6 selects 24A siRNAs mainly via a strong 5′ adenine preference. Domain swapping experiments showed that the coordination of the MID and PIWI domains, but not the previously known nucleotide specificity loop in the MID domain, determines the 5′ A preference of AGO4 and AGO6. Our study suggests that accumulated mutations in the PIWI and MID domains of AGO4 clade proteins conferred distinct small RNA selectivity to these AGOs after gene duplication, which may contribute to elaborate epigenetic regulation in plants.

## MATERIAL AND METHODS

### 1. Bioinformatics re-analysis of Pol IV-derived RNAs and siRNAs

#### 1.1 Defining P4RNAs

The already published PATH (parallel analysis of tail and head) library (SRR2000577(*dcl2/3/4* deficient mutant)) was downloaded from Sequence Read Archive. After the TruSeq small RNA adapter sequence was trimmed by cutadapt (26), the sequence library was mapped to the previously defined Pol IV clusters (including both SHH1-dependent and SHH1-independent case (27)) by bowtie (28). P4RNAs sharing a 5′ end were combined into one, and P4RNAs >30 nt in length and >50 reads were used in following analysis. 8488 kinds of P4RNAs were defined.

#### 1.2 Mapping AGO-bound small RNAs to defined P4RNAs

After the TruSeq small RNA adapter sequence was trimmed by cutadapt, AGO4-bound RNAs (29) (SRR189808, AGO4 wild-type) and AGO6-bound RNAs (30) (SRR1266014, AGO6 wild-type) were mapped to the defined P4RNAs by bowtie. To ensure that the siRNAs were derived from the Pol IV-transcripts, only the siRNAs whose 5′ ends were exactly aligned with P4R2 RNAs were selected for further processing.

#### 1.3 Grouping P4RNAs by conversion efficiency (CE)

The conversion efficiency (CE) was calculated by dividing the number of reads of AGO4/6-bound small RNAs by the number of reads of P4R2 RNAs. The P4R2 RNAs were evenly divided into four bins in ascending order of CE value, from the lowest bin 1 to the highest bin 4.

### 2. Plasmid construction

The primers used in this study are listed in **Supplementary Table S1**.

#### pBYL-AtAGO6

A DNA fragment containing the AGO6 CDS was amplified by PCR from pCRHA-AGO6 (11) using oligo No. 1 and oligo No. 2, digested with AscI, and inserted into the corresponding region of pBYL2 (31).

#### pBYL-AtAGO4

A DNA fragment containing the AGO4 CDS was amplified by PCR from pCRHA-AGO4 (11) using oligo No. 3 and oligo No. 4, digested with AscI, and inserted into the corresponding region of pBYL2 (31).

#### pBYL-3×FLAG-SUMO-AtAGO4

Two DNA fragments were prepared by PCR: the 3×FLAG-SUMO fragment was amplified from pBYL-3×FLAG -SUMO-AtAGO1 (32) with oligo No. 5 and oligo No. 6, and the AGO4 CDS was amplified from pBYL-AtAGO4 using oligo No. 7 and oligo No. 4. These fragments were further amplified with oligo No. 5 and oligo No. 4, digested with AscI, and inserted into the corresponding region of pBYL2 (31).

#### pBYL-3×FLAG-SUMO-AtAGO6

A DNA fragment containing the 3×FLAG-SUMO was amplified by PCR from pBYL-3×FLAG-SUMO-AtAGO4 with oligo No. 5 and oligo No. 8. Another DNA fragment containing the AtAGO6 CDS was amplified from pBYL-AtAGO6 using oligo No. 9 and oligo No. 2. These fragments were further amplified with oligo No. 2 and oligo No. 5, digested with AscI, and inserted into the corresponding region of pBYL2.

#### pBYL-3×FLAG-AtAGO4

A DNA fragment containing 3×FLAG was amplified from pBYL-3×FLAG-SUMO-AtAGO4 with oligo No. 10 and oligo No. 11. Another DNA fragment containing the AtAGO4 CDS was also amplified from pBYL-3×FLAG-SUMO_AtAGO4 with oligo No. 12 and oligo No. 4. These fragments were further amplified with oligo No. 10 and oligo No. 4, digested with AscI, and inserted into the corresponding region of pBYL2.

#### pBYL-3×FLAG-AtAGO6

A DNA fragment containing the 3×FLAG was amplified from pBYL-3×FLAG-SUMO-AtAGO6 with oligo No. 10 and oligo No. 13. Another DNA fragment containing the AtAGO6 CDS was also amplified from 3×FLAG-SUMO-AtAGO6 with oligo No. 14 and oligo No. 2. These fragments were further amplified with oligo No. 10 and oligo No. 2, digested with AscI, and inserted into the corresponding region of pBYL2.

#### pBYL-3×FLAG-AGO4L1

A DNA fragment corresponding to the first half of 3×FLAG-AGO4-CDS (926–2790 nt) was amplified from pBYL-3×FLAG-AGO4 using oligo No. 15 and oligo No. 16. Another DNA fragment corresponding to the second half of the AGO4 CDS (2758–3819 nt) was amplified from pBYL-3×FLAG-AGO4 using oligo No. 17 and oligo No. 18. These two fragments were mixed and assembled into pBYL2 vector using the NEBuilderHiFi Assembly Kit (NEB).

#### pBYL-3×FLAG-AGO4L6

A DNA fragment corresponding to the first half of 3×FLAG-AGO4 CDS (926–2784 nt) was amplified from pBYL-3×FLAG-AGO4 using oligo No. 15 and oligo No. 19. Another DNA fragment corresponding to the second half of the AGO4 CDS (2767-3822 nt) was amplified from pBYL-3×FLAG-AGO4 using oligo No. 18 and oligo No. 20. These two fragments were mixed and assembled into pBYL2 vector using the NEBuilderHiFi Assembly Kit (NEB).

#### pBYL-3×FLAG-AGO6L1

A DNA fragment corresponding to the first half of 3×FLAG-AGO6 CDS (926–2673 nt) was amplified from pBYL-3×FLAG-AGO6 using oligo No. 15 and oligo No. 21. Another DNA fragment corresponding to the second half of the AGO6 CDS (2656–3681 nt) was amplified from pBYL-3×FLAG-AGO6 using oligo No. 22 and oligo No. 23. These two fragments were mixed and assembled into pBYL2 vector using the NEBuilderHiFi Assembly Kit (NEB).

#### pBYL-3×FLAG-AGO6L4

A DNA fragment corresponding to the first half of 3×FLAG-AGO6 CDS (926–2676 nt) was amplified from pBYL-3×FLAG-AGO6 using oligo No. 15 and oligo No. 24. Another DNA fragment corresponding to the second half of the AGO6 CDS (2659–3684 nt) was amplified from pBYL-3×FLAG-AGO6 using oligo No. 23 and oligo No. 25. These two fragments were mixed and assembled into pBYL2 vector using the NEBuilder HiFi Assembly Kit (NEB).

#### pBYL-3×FLAG-AGO4M6

A DNA fragment corresponding to the MID domain of AtAGO6 (2518–2916 nt) was amplified from pBYL-3×FLAG-AGO6 using oligo No. 26 and oligo No. 27. Another DNA fragment was amplified from pBYL-3×FLAG-AGO4 using oligo No. 28 and oligo No. 29. These two fragments were mixed and assembled using the NEBuilder HiFi Assembly Kit (NEB).

#### pBYL-3×FLAG-AGO4P6

A DNA fragment corresponding to the PIWI domain of AtAGO6 (2914–3765 nt) was amplified from pBYL-3×FLAG-AGO6 using oligo No. 30 and oligo No. 31. Another DNA fragment was amplified from pBYL-3×FLAG-AGO4 using oligo No. 32 and oligo No. 33. These two fragments were mixed and assembled using the NEBuilder HiFi Assembly Kit (NEB).

#### pBYL-3×FLAG-AGO6M4

A DNA fragment corresponding to the MID domain of AtAGO4 (2416–2805 nt) was amplified from pBYL-3×FLAG-AGO4 using oligo No. 34 and oligo No. 35. Another DNA fragment was amplified from pBYL-3×FLAG-AGO6 using oligo No. 36 and oligo No. 37. These two fragments were mixed and assembled using the NEBuilder HiFi Assembly Kit (NEB).

#### pBYL-3×FLAG-AGO6P4

A DNA fragment corresponding to the PIWI domain of AtAGO4 (2809–3681 nt) was amplified from pBYL-3×FLAG-AGO4 using oligo No. 31 and oligo No. 38. Another DNA fragment was amplified from pBYL-3×FLAG-AGO6 using oligo No. 32 and oligo No. 39. These two fragments were mixed and assembled using the NEBuilder HiFi Assembly Kit (NEB).

### 3. Preparation of AGO mRNAs

mRNAs were *in vitro* transcribed from NotI-digested (NEB) plasmids using the AmpliScribe T7 High Yield Transcription Kit (Lucigen), followed by capping with Script-Cap m^7^G Capping System (Cell Script). Poly(A)-tails were added to the transcripts using the A-Plus Poly(A) Polymerase Tailing Kit (Cell Script).

### 4. Preparation of BY-2 cell lysate

The BY-2 lysate was prepared as described previously (33).

### 5. Preparation of radiolabeled double-stranded small RNA

The sequences of the guide and passenger RNAs used in this study are shown in **Supplementary Table 2**. Single-stranded RNAs with a 5′ hydroxyl group (OH) were synthesized by GeneDesign Inc. (Osaka Japan). The guide strand was radiolabeled using T4 PNK and [γ-32P] ATP. The passenger strand with a 5′ hydroxyl group was phosphorylated using T4 PNK and ATP. The guide and passenger strands (with 5′ monophosphate or C6-amino linker) were heat-annealed in lysis buffer at 95°C for 2 min and 25°C for 30 min. The size of the annealed RNA and guide strand were checked by 12% polyacrylamide native gel containing 1 mM Mg^2+^.

### 6. *In vitro* RISC assembly assay

Thirty microliters of BY-2 lysate, 15 μl of substrate mixture (3 mM ATP, 0.4 mM GTP, 100 mM creatine phosphate, 200 μM each of 20 amino acids, 320 μM spermine, 0.4 U/μl creatine phosphokinase (Calbiochem)) and 3 μl of 1 μg/μl AGO mRNA were incubated for 90 min at 25°C. Translation of AGO mRNAs was terminated by adding 2.4 μl of 20 mM cycloheximide. Nine-microliter aliquots were incubated with 1 μl of 50 nM radiolabeled small RNA duplexes at 25°C for 30 min. The reaction was mixed with anti-FLAG antibody (Sigma) immobilized on 2.5 μl of Dynabeads protein G (Invitrogen) and incubated for 30 min on ice. After incubation, the beads were washed three times with 500 μl of lysis buffer [30 mM HEPES (pH 7.4), 100 mM KOAc and 2 mM Mg(OAc)_2_] containing 1% Triton X-100 and 800 mM NaCl, and treated with proteinase K. After ethanol precipitation, the samples were resuspended in 10 μl of 2x formamide dye [10 mM EDTA pH 8.0, 98% (wt/vol) deionized formamide, 0.025% (wt/vol) xylene cyanol, and 0.025% bromophenol blue], and 5 μl aliquots were separated on an 15% denaturing acrylamide gel. Gels were analyzed by Typhoon FLA 7000 image analyzer (GE Healthcare) and quantified using MultiGauge software (Fujifilm Life sciences). Graphs were prepared using GraphPad Prism 9. In Supplementary Figure 2, before adding small RNA duplexes, geldanamycin (GA) at a final concentration of 1 mM and DMSO at a final concentration of 1% were added to the experimental and control group, respectively. Equimolar amounts of non-radiolabeled small RNA duplexes were used for western blotting of immunoprecipitated AGO proteins in Figure 2B and 2D.

**Figure 2.**
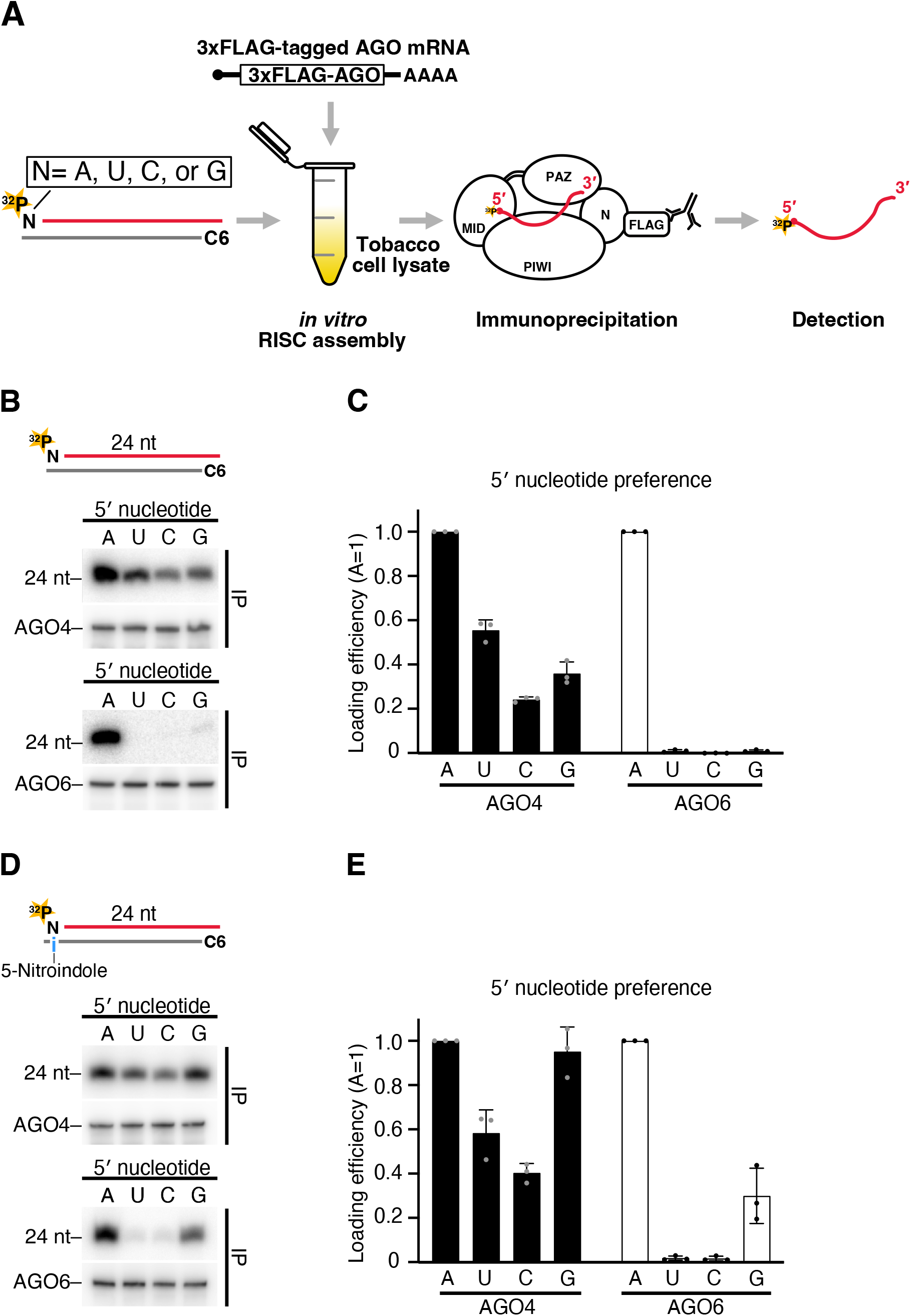
AGO4 and AGO6 actively select 24A siRNAs via distinct mechanisms. (A) Schematic of RISC assembly *in vitro*. To form RISC, 3×FLAG-tagged AGO mRNAs were translated in tobacco cell lysate, then incubated with radiolabeled small RNA duplexes containing specific 5′ nucleotides. Co-immunoprecipitation was performed using anti-FLAG coated beads to purify AGO4/6-bound siRNAs. The radioactivity was quantified after separation by a denaturing UREA PAGE. (B) AGO4/6–RISC assembly using siRNA duplexes bearing different 5′ nucleotide on the guide strand. AGO4 and AGO6 actively select 24A siRNAs. (Top) Schematic for siRNA duplexes used in this study. The guide strand was 5′ radiolabeled. The passenger strand has an amino C6 linker at the 5′ end that prevents the passenger strand from being loaded onto AGO4/6. (Middle) Small RNAs loaded into AGO4 (upper panel) and western blotting of AGO4 proteins expressed in the tobacco BY-2 lysate after immunoprecipitation with anti-FLAG antibody (lower panel). (Bottom) Small RNAs loaded into AGO6 and western blotting of AGO6 proteins expressed in the tobacco BY-2 lysate after immunoprecipitation with anti-FLAG antibody. (C and E) Quantification of loaded siRNAs in (B) and (D), respectively. The band intensity of siRNAs was normalized to the value of 5′ A. The graphs show the mean ± SD from three technically independent experiments. Loading efficiency of each experiment of AGO4 and AGO6 was shown as gray and black dots. (D) 5′ nucleotide preference of AGO4 and AGO6 when thermodynamic stability and nucleotide identity are separated. AGO4 accepted all four siRNAs with a slight preference for the siRNAs with purine nucleotides, whereas siRNAs with 5′ A were still predominantly incorporated into AGO6. (Top) 5-nitroindole (i) was introduced at the nucleotide position of the passenger strand facing the 5′ terminal nucleotide of the guide strand. See also (B).

## RESULTS

### P4RNAs with 5′ A and/or unstable ends are successfully converted into mature siRNAs

In general, the small RNA preference of a particular AGO can be estimated by comparing the sequences of small RNAs co-immunoprecipitated (co-IPed) with the AGO and those of total (input) small RNAs. For example, the small RNAs co-IPed with AGO1, AGO2, and AGO5 contain a larger proportion of those with U, A, and C at their 5′ ends than total small RNAs, suggesting that AGO1, 2, and 5 have 5′ U, A, and C selectivity, respectively (10–12). Previous studies showed that AGO4 and AGO6 associate with 24-nt small RNAs with 5′ A (18). However, since more than 60% of the 24-nt small RNAs in plants naturally have adenine at the 5′ end (18), it is challenging to accurately analyze the 5′ nucleotide selectivity of AGO4 and AGO6 by simply comparing the total and co-IPed small RNAs. Instead, we calculated the conversion efficiency (CE) by dividing the number of reads of AGO4/6-bound small RNAs by the number of reads of precursor RNAs (P4RNAs) that share the same sequence with the small RNAs. This value indicates the efficiency with which a given precursor RNA sequence is converted into AGO-bound mature small RNAs, reflecting both dicing and loading efficiency. Higher values indicate a tendency to form RISC more efficiently, while lower values indicate a tendency to form RISC less efficiently. We obtained P4RNAs accumulated in a dicing deficient mutant *Arabidopsis thaliana* (*dcl2/3/4*) and siRNAs co-IPed with AGO4/6 in wild-type plants from a public database (21, 27, 29, 30). Then, we mapped the siRNAs to the P4RNAs (see also materials and methods). We divided the P4RNAs evenly into four bins in ascending order of CE value, from the lowest bin 1 to the highest bin 4 (Figure 1B and C). P4RNAs with higher CE tended to have a stronger 5′ A bias in both AGO4 and AGO6 (Figure1D and E), suggesting that P4RNAs with 5′ A are efficiently processed and loaded into AGO4 and AGO6 *in vivo*.

The thermodynamic stability of the base pairing at the end of small RNA duplexes can affect the strand selection of small RNAs in animals (34, 35). Therefore, we wondered if the thermodynamic stability also affects the small RNA loading efficiency in AGO4 and AGO6. To separate it from the effect of 5′-nucleotide identity, we investigated the nucleotide distribution at the second position from the 5′ end (g2 position) of the small RNAs with 5′ A. P4RNAs with higher CE tended to have an A/U nucleotide rather than a C/G nucleotide at the g2 position (Figure 1F and G). We obtained similar results when we analyzed 5′ G P4RNAs (Supplementary Figure 1A and B). Thus, the thermodynamic stability at the 5′ end also affects the conversion from P4RNAs to AGO4/6-bound mature small RNAs. Taken together, we conclude that both the 5′ A nucleotide and the thermodynamically unstable end promote the conversion from P4RNAs to AGO4/6-bound mature small RNAs *in vivo*.

### AGO4 and AGO6 actively select 24A siRNAs via distinct mechanisms

Previous reports showed that DCL3 prefers to cleave dsRNA precursors from the end with 5′ A and/or unstable sequence (23–25). Thus, we cannot exclude the possibility that the siRNAs with the 5′ A and the thermodynamically unstable ends are enriched at the DCL3-mediated cleavage steps but not at the loading step. To solve this problem, we applied a cell-free system derived from tobacco BY-2 protoplasts, which faithfully recapitulates the RISC assembly processes (4, 32, 36, 37). We first *in vitro* translated mRNAs encoding 3×FLAG-tagged AGO4 or AGO6 ORFs in the cell-free system. To assemble RISC, we then incubated 24-nt siRNA duplexes, of which one strand was radiolabeled with ^32^P, while the other strand had a 5′ C6 amino linker that blocks loading of this strand as the guide strand. Finally, we IPed 3×FLAG-tagged AGO4/6 with a FLAG antibody, and separated the co-IPed siRNAs in a denaturing gel (Figure 2A). We successfully detected siRNAs co-immunoprecipitated with AGO4 and AGO6, whereas HSP90-inhibitor geldanamycin (GA) abrogated loading of siRNAs (Supplementary Figure 2A–C). Thus, our system faithfully recapitulates HSP90 chaperone-dependent 24-nt siRNA loading.

We next investigated the loading efficiencies of 24-nt siRNAs with different nucleotides (A, U, C and G) at the 5′ end. The 24A siRNAs were most efficiently loaded into both AGO4 and AGO6 (Figure 2B and C). Thus, AGO4 and AGO6 actively select 24A siRNAs at the loading step. However, the degree of specificity for 24A siRNAs differed between the two AGOs: AGO6 specifically loaded 24A siRNAs, whereas AGO4 could also load 24U, 24C and 24G siRNAs with an efficiency of ∼50%, ∼20% and ∼30%, respectively, compared to 24A siRNAs (Figure 2B and C).

To separate the effects of nucleotide specificity and thermodynamic stability, we introduced 5-nitroindole, a universal base that hybridizes with all four bases with similar affinities (38, 39), at the position on the passenger strand facing the 5′ end of the guide strand (Figure 2D). AGO4 accepted all four siRNAs with a slight preference for the siRNAs with purine nucleotides (Figure 2D and E) at the 5′ end. Thus, not only the nucleotide identity but also the thermodynamic stability at the 5′ end of siRNAs plays critical roles in selecting the 24A siRNAs in AGO4. Although the use of the passenger strand with 5-nitroindole slightly enhanced loading of 24G siRNAs into AGO6, 24A siRNAs were still predominantly incorporated into AGO6 (Figure 2D and E). Taken together, we concluded that AGO4 and AGO6 actively select 24A siRNAs via different mechanisms: AGO4 selects 24A siRNAs via a combination of a 5′ purine preference and a preference for unstable base-pairing at the 5′ end, whereas AGO6 selects 24A siRNAs mainly via a strong 5′ A preference.

### AGO6 forms RISC more efficiently with shorter siRNAs than AGO4

A previous study showed that AGO6 can load not only 24A siRNAs but also 21-nt siRNAs with various 5′ nucleotides in both wild-type Col-0 and *ddm1* mutant plants, in which TEs are transcriptionally active (30). To determine how efficiently AGO4 and AGO6 can load 21-nt siRNAs, we performed *in vitro* RISC assembly with 21-nt and 24-nt siRNAs and their mixture. AGO4 preferentially bound 24-nt siRNAs compared to 21-nt siRNAs, whereas AGO6 was able to load both (Figure 3A and B). To elucidate the precise range of siRNA lengths to which AGO4 and AGO6 can bind, we performed *in vitro* RISC assembly using an oligo pool containing equal amounts of siRNA duplexes ranging from 19 to 25 nucleotides (Figure 3A). Both AGO4 and AGO6 preferentially bound to long (23–25 nt) siRNA, with the highest efficiency at 24-nt siRNAs (Figure 3C and D). AGO6 also bound short (21–22 nt) siRNAs compared to AGO4 (Figure 3C and D), confirming the previous notion *in vivo* (30).

**Figure 3.**
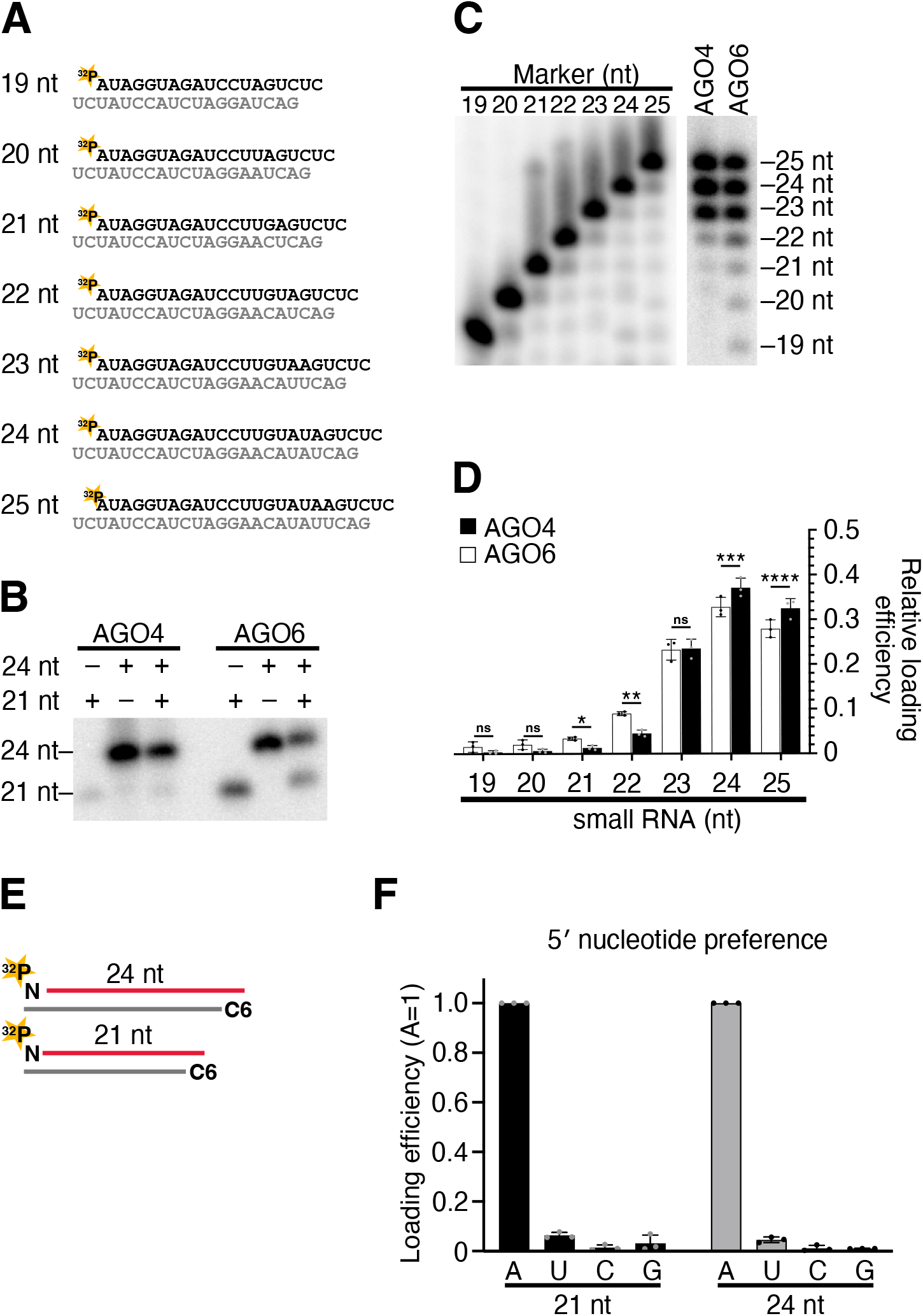
AGO6 forms RISC more efficiently with shorter siRNAs than AGO4. (A) The 19–25-nt siRNA duplexes used in this study. The 5′ end of the guide strand was radiolabeled with ^32^P. (B) *In vitro* AGO4/6-RISC assembly with 21- and 24-nt siRNAs. AGO4 preferentially bound 24-nt siRNAs compared to 21-nt siRNAs, whereas AGO6 was able to load both. (C) *In vitro* AGO4/6-RISC assembly using a mixture of 19–25-nt siRNA duplexes. Although both AGO4 and AGO6 preferentially bound to long (23–25 nt) siRNAs, AGO6 also bound short (21–22 nt) siRNAs compared to AGO4. (D) Quantification of loaded siRNAs in (C). The relative band intensity of each length of siRNA was calculated with the total band intensity as 1. The graphs show the mean ± SD from three technically independent experiments (AGO4, black dots; AGO6, gray dots). Benjamini–Hochberg procedure (false discovery rate approach)-corrected p values from multiple paired t tests are as follows: *p = 0.029415, **p = 0.029415, ***p = 0.006793, ****p = 0.048948. ns, not significant. (E) AGO6-RISC assembly using 21-nt and 24-nt siRNA duplexes bearing different 5′ nucleotide on the guide strand. The guide strand was 5′ radiolabeled. The passenger strand has an amino C6 linker at the 5′ end that prevents the passenger strand from being loaded onto AGO6. AGO6 strictly selected siRNAs with 5′ A, regardless of the siRNA length. (F) Quantification of loaded siRNAs in (E). The band intensity of siRNAs was normalized to the value of 5′ A. The graphs show the mean ± SD from three technically independent experiments.

We next asked if the 5′ nucleotide bias can be changed when 21-nt, instead of 24-nt, siRNAs are loaded into AGO6. We performed *in vitro* RISC assembly experiments with 21-nt siRNA duplexes, of which one strand was radiolabeled with ^32^P, while the other strand had a 5′ C6 amino linker to block loading (Figure 3E). As in the case with 24-nt siRNAs, AGO6 strictly selected 21-nt siRNAs with 5′ A (Figure 3F). Thus, small RNA length does not change the strict 5′ A preference of AGO6 *in vitro*.

### The nucleotide specificity loop in the MID domain does not determine the 5′ -nucleotide preference of AGO4 and AGO6

Next, we asked how AGO4 and AGO6 recognize the 5′ nucleotide. It has been suggested that the nucleotide specificity loop in the MID domain plays an important role in 5′-nucleotide selection in hAGO2 and AtAGO1 (9, 13) (Figure 4A). To test if the loops also determine the degree of 5′ A preference of AGO4 and AGO6, we performed *in vitro* RISC assembly using loop-swapped chimeric AGO4 and AGO6 (AGO4L6 and AGO6L4, Figure 4B) and siRNA duplexes with 5-nitroindole on the passenger strand (Figure 4C). Strikingly, AGO4L6 and AGO6L4 showed the same 5′-nucleotide preference as original AGO4 and AGO6 (Figure 4D and E). Thus, the loop structures in AGO4 and AGO6 do not determine the strictness of 5′ A recognition in the RISC assembly. To further investigate the function of the nucleotide specificity loop in the 5′-nucleotide recognition, we replaced the loop of AGO4 and AGO6 with that of AGO1, which prefers 5′ U (Supplementary Figure 3A). Similarly, the loop switching barely changed their 5′-nucleotide preference (Supplementary Figure 3B and C). Based on these results, we conclude that the nucleotide specificity loop does not account for the 5′-nucleotide selection in AGO4 and AGO6 in *Arabidopsis*.

**Figure.4.**
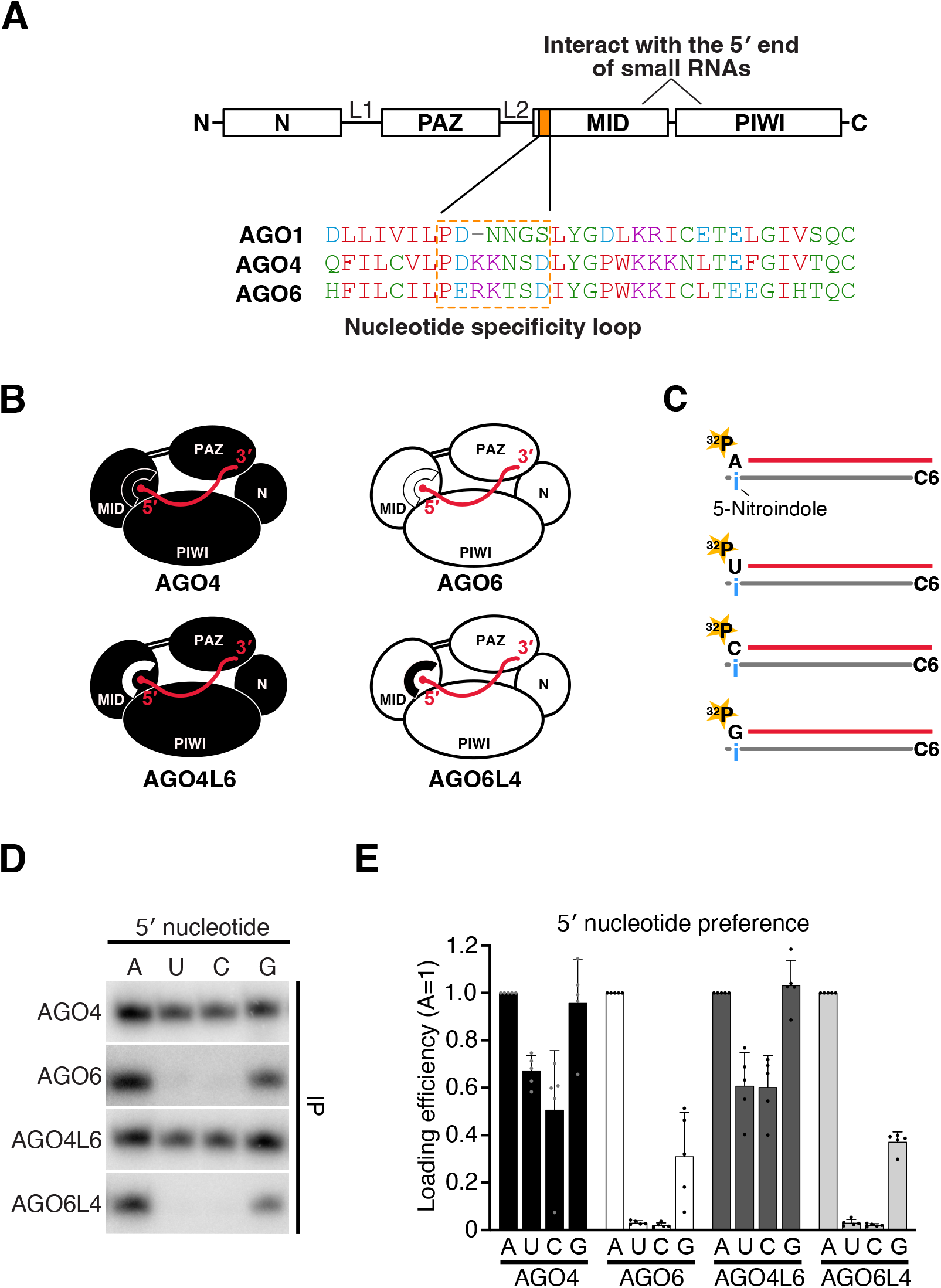
The nucleotide specificity loop in the MID domain does not determine the 5′-nucleotide preference of AGO4 and AGO6. (A) Schematic domain architecture of Argonaute protein. The 5′ end of small RNA interacts with the 5′ end binding pocket formed by the MID and PIWI domains. The nucleotide specificity loop is in the MID domain. The sequences of nucleotide specificity loops of AGO1, AGO4 and AGO6 are shown inside the orange frame. (B) Schematic of chimeric AGO proteins whereby the nucleotide-specific loops of AGO4 and AGO6 were swapped. (C) Schematic for siRNA duplexes used in this study. The guide strand was 5′ radiolabeled. The passenger strand has an amino C6 linker at the 5′ end that prevents the passenger strand from being loaded onto AGO. 5-nitroindole (i) was introduced at the nucleotide position of the passenger strand facing the 5′ terminal nucleotide of the guide strand. (D) *In vitro* RISC assembly with AGO4/6 chimeric proteins. AGO4L6, like AGO4, accepted all four siRNAs with a slight 5′ purine preference. AGO6L4 predominantly showed a 5′ A preference as in AGO6. (E) Quantification of loaded siRNAs in (D). The band intensity of siRNAs was normalized to the value of 5′ A. The graphs show the mean ± SD from five technically independent experiments. Note that the experiments with chimeric AGO4/6s in Figures 4E, 5C, and 5E were performed simultaneously. For the experimental controls, wild-type AGO4 and AGO6, the samples were electrophoresed in three different gels corresponding to Figures 4E, 5C, and 5E, resulting in similar quantitative results in those Figures.

### Both MID and PIWI domains are involved in 5′ nucleotide selection

We next swapped the whole MID domain between AGO4 and AGO6 (Figure 5A), and performed RISC assembly assay using siRNA duplexes with 5-nitroindole on the passenger strand (Figure 4C). When the MID domain of AGO4 was replaced with that of AGO6 (hereafter named AGO4M6), the 5′ U and 5′ C preference significantly decreased compared to the wild-type AGO4 (Figure 5B and C). However, the 5′ G preference barely changed (Figure 5B and C). On the other hand, the counterpart chimera, AGO6M4, showed no changes in 5′-nucleotide preference compared to AGO6 (Figure 5B and C). These results suggest that 1) the MID domain is partially involved in 5′-nucleotide selection in AGO4, and 2) domains other than the MID domain are also involved in 5′-nucleotide selection in both AGO4 and AGO6.

**Figure 5.**
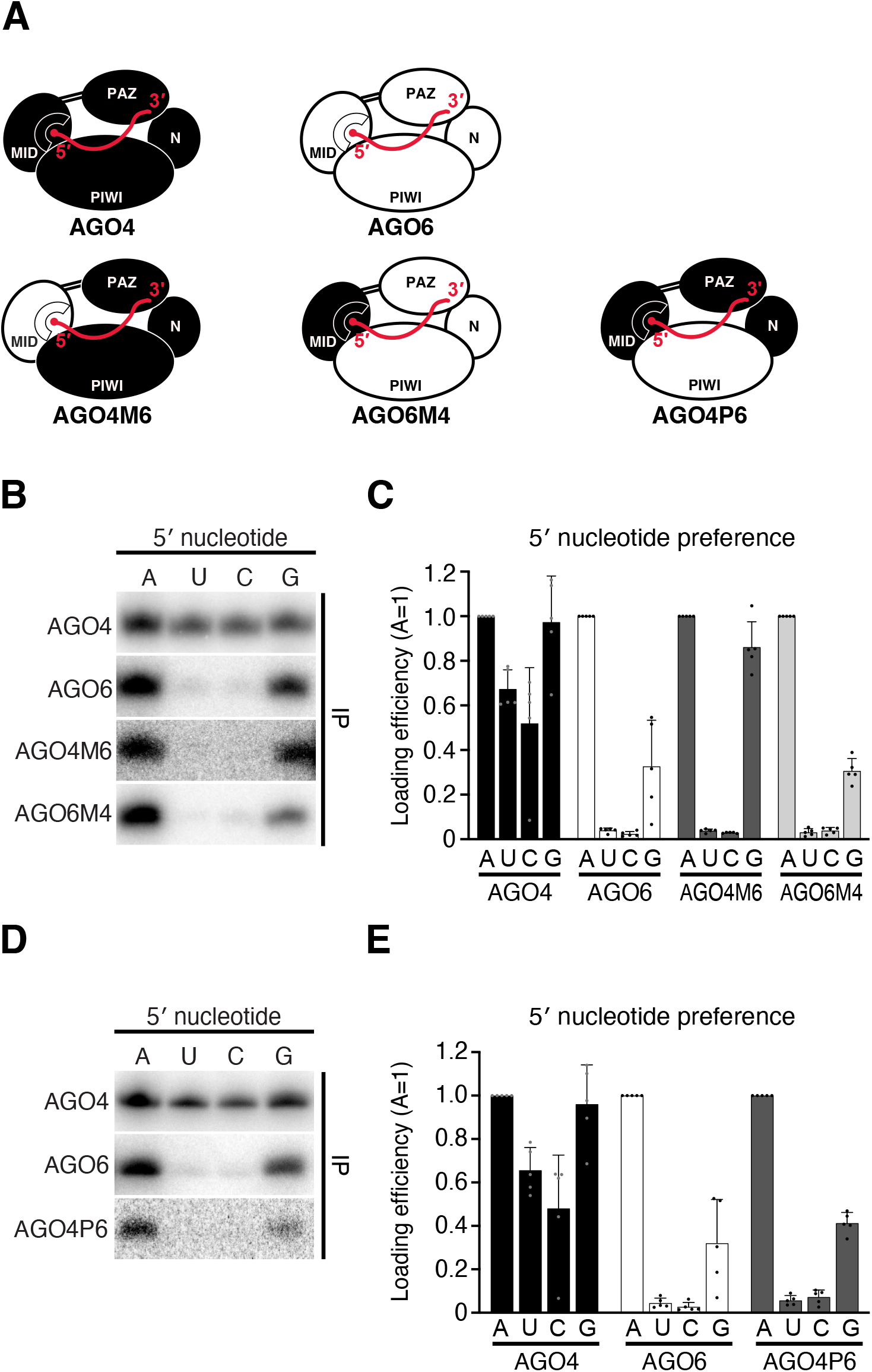
Both MID and PIWI domains are involved in 5′ nucleotide selection. (A) Schematic diagram of chimeric proteins with MID or PIWI domains swapped between AGO4 and AGO6. (B) *In vitro* RISC assembly with AGO4/6 chimeric proteins. AGO4M6 showed a predominant preference for 5′ A and 5′ G. AGO6M4 showed a predominant 5′ A preference as in AGO6. (C) Quantification of loaded siRNAs in (B). The band intensity of siRNAs was normalized to the value of 5′ A. The graphs show the mean ± SD from five technically independent experiments. See also Figure 4E. (D) *In vitro* RISC assembly with a AGO4/6 chimeric protein. AGO4P6 showed a predominant 5′ A preference as in AGO6. (E) Quantification of loaded siRNAs in (D). The band intensity of siRNAs was normalized to the value of 5′ A. The graphs show the mean ± SD from five technically independent experiments. See also Figure 4E.

The above domain swapping experiment suggests that the MID domain alone does not determine the selectivity of the 5′ nucleotide. Since the 5′-nucleotide binding pocket is located between the MID domain and the PIWI domain, it is possible that the PIWI domain is also involved in the selection of the 5′-nucleotide. To test this hypothesis, we swapped the PIWI domain between AGO4 and AGO6 (Figure 5A). Unfortunately, AGO6P4, which is AGO6 chimera with the PIWI domain of AGO4, showed no small RNA binding activity, presumably due to folding failure (data not shown). However, simply replacing the PIWI domain of AGO4 with that of AGO6 resulted in the same strict 5′ A nucleotide selectivity as AGO6 (AGO4P6, Figure 5D and E). Overall, our results suggest that both the MID and PIWI domains of AGO4 are required for the loose 5′ A selectivity of AGO4, and that the PIWI domain of AGO6 plays a critical role in the strong 5′ A selectivity of AGO6.

## DISCUSSION

### AGO4 and AGO6 select siRNAs with different characteristics via distinct mechanisms

Previous studies have shown that AGOs select small RNAs by sensing the 5′ nucleotide identity and the thermodynamic stability at the 5′ terminal base pair (10, 11, 34, 35). In this study, we found that their contributions for small RNA selection differ between the two AGO4 clade proteins, AGO4 and AGO6. AGO6 mainly selects the 5′ A siRNAs by recognizing the nucleotide identity, while AGO4 preferentially loads siRNAs with a weak 5′ terminal base pair (Figure 6A). In addition to the differences in the mechanisms for siRNA selection, AGO4 and AGO6 showed different degrees of preference for 5′ A siRNAs, with AGO6 showing a stronger preference for 5′ A than AGO4. These *in vitro* results are consistent with the previous *in vivo* small RNA-IP analyses, in which 5′ A siRNAs were more enriched in the AGO6-IP fraction than in the AGO4-IP fraction (18). Interestingly, in contrast to the strictness of 5′ nucleotide recognition, AGO6 showed a more flexible length selectivity than AGO4 (Figure 3B-D); AGO4 preferentially loaded long siRNAs (23-25 nt), while AGO6 loaded not only long siRNAs but also short siRNAs (19-22 nt) (Figure 3B and 3C). Thus, AGO4 and AGO6 may have changed the repertoire of their binding siRNAs during divergence after gene duplication.

**Figure 6.**
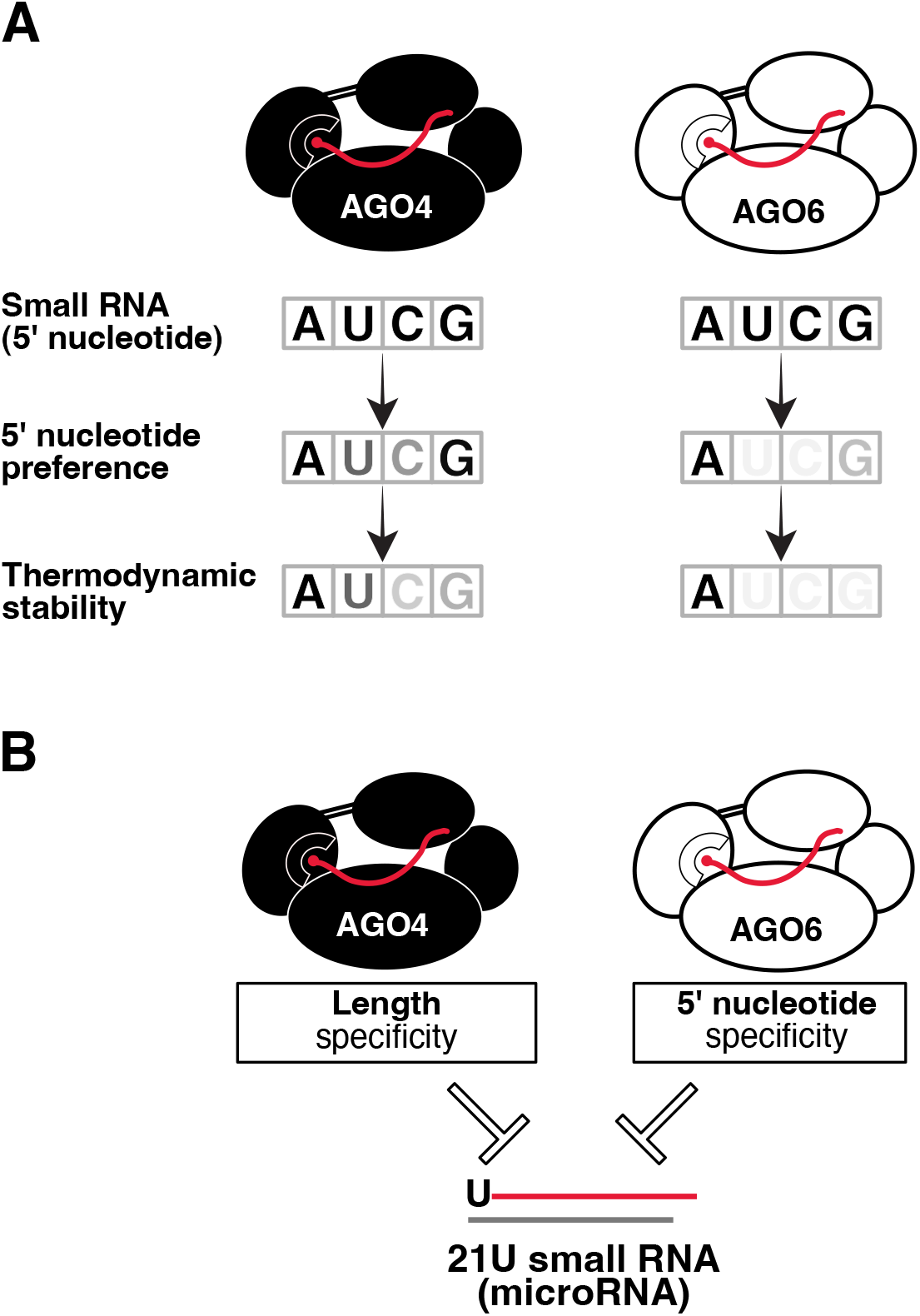
AGO4 and AGO6 have distinct small RNA selection mechanisms. (A) Schematic of distinct 5′ A selection mechanisms of AGO4 and AGO6. AGO6 mainly selects the 5′ A siRNAs by recognizing the nucleotide identity, while AGO4 loads 5′ A sRNAs through a combination of 5′ purine preference and preference for weak 5′ terminal base pair. (B) A model for 21U miRNA rejection by AGO4 and AGO6. Because of the property of AGO4 to bind only 23-25 nt long siRNAs and AGO6 to form RISC with only 5′ A siRNAs, these AGOs are unable to bind 21U miRNAs, avoiding unwanted transcriptional repression of the miRNA locus.

### PIWI and MID domains of plant AGOs are involved in 5′ nucleotide selectivity

Until now, it was thought that the nucleotide specificity loop in the MID domain is the determinant of small RNA sorting in plants. This model was proposed based on NMR titration and structural analyses with the MID domain and nucleoside monophosphates (9, 13). However, our *in vitro* RISC assembly using loop-switched chimeric AGOs showed that the nucleotide specificity loop is not the determinant for small RNA sorting in AGO4 and AGO6 (Figure 4 and Supplementary Figure 3). Similar results were obtained in experiments with PIWI clade proteins, one of the Argonaute families that represses TEs in germline cells in animals: mutation or deletion of the nucleotide specificity loop in *Drosophila* PIWI did not affect the preference for 5′ U PIWI-interacting RNAs (piRNAs) (40). These results suggest that structures other than nucleotide specificity loops contribute to the 5′ nucleotide preference in both AGO and PIWI proteins.

Our domain swapping experiments showed that replacing the MID or PIWI domains of AGO4 with those of AGO6 switches the loose 5′ A selectivity of AGO4 to AGO6-like strong 5′ A selectivity (Figure 5). Interestingly, replacing the PIWI domain of AGO4 with that of AGO6 perfectly mimicked AGO6’s 5′ nucleotide preference (Figure 5D and 5E). These results suggest that not only the MID domain, but also the PIWI domain plays important roles in small RNA sorting. In *Rhodobacter sphaeroides* AGO (RsAgo), a bacterial AGO possessing a 5′ U preference, the side chain of R754 of the PIWI domain forms hydrogen bonds with O4 of the 5′ U of the guide RNA strand (41). The PIWI domain in eukaryotic AGOs may also be involved in 5′-nucleotide selection by directly interacting with the 5′ nucleotide. Alternatively, the difference in the size of the pocket space formed by the MID and PIWI domains may affect the selectivity of the 5′ nucleotides of AGO4 and AGO6. Future structural analysis of the complex consisting of a full-length AGO and a small RNA will elucidate the detailed mechanism of 5′ nucleotide selection in AGO4/6-RISCs.

### Biological significance of distinct small RNA selection strategies of AGO4 and AGO6

AGO4 clade proteins function in two RdDM pathways: the Pol IV-RDR2-mediated canonical RdDM pathway and the Pol II-RDR6-mediated non-canonical RdDM pathway. In the canonical pathway, Pol IV, RDR2, and DCL3 act sequentially to generate 24-nt siRNAs whose 5′ terminal nucleotide is predominantly adenine. After forming RISC with AGO4/6, the guide strand of siRNA directs the RISC to Pol V-nascent transcripts, inducing DNA methylation of target gene via DOMAINS REARRANGED METHYLTRANSFERASE 2 (DRM2). This process is considered as a “self-reinforcing” pathway, because Pol IV and Pol V are recruited to the DNA loci modified by RdDM. In this study, we found that both AGO4 and AGO6 actively select 5′ A siRNAs, which are enriched in the biogenesis pathway. These results suggest that the AGO4 clade proteins have evolved their substrate specificity to induce efficient canonical RdDM.

The non-canonical RdDM pathway functions to establish epigenetic silencing of reactivated TEs and induces DNA methylation at non-TE loci. In this pathway, Pol II transcripts from actively expressed TEs or viruses are first cleaved by AGO1-RISC and converted into dsRNA by RDR6, which are then diced into 21–24 nt siRNAs by DCL1/2/3/4 (42). These siRNAs with different lengths and 5′ nucleotides are loaded into AGO4 clades proteins to induce DNA methylation (42, 43). Our study shows that AGO4 accepts not only 24A siRNAs, but also 24-nt siRNAs with other nucleotides at the 5′ end. This loose 5′ nucleotide selectivity may allow AGO4 to accept 24-nt siRNAs with various 5′ nucleotides generated in the non-canonical pathway. On the other hand, AGO6 can form RISC not only with 24-nt siRNAs but also with shorter (19–22 nt) siRNAs. This property of AGO6 may allow RdDM to be triggered with 21–22 nt siRNAs that are processed by DCL4 and DCL2. In fact, previous studies have reported that in *dcl3/ddm1* mutant *Arabidopsis* (TE-activated mutants), AGO6 forms RISCs with short siRNAs to trigger RdDM at the activated TE loci (30). Importantly, the property of AGO4 to bind only 24–25 nt long siRNAs and the property of AGO6 to form RISC only with 5′ A siRNAs prevent those AGOs from binding to 21U miRNAs, avoiding unwanted transcriptional repression of miRNA loci (Figure 6B). Thus, the distinct small RNA selection mechanisms of AGO4 and AOG6 may help to achieve robust epigenetic regulation at target loci such as TEs, without affecting the regulation of normal gene expression.

## AUTHOR CONTRIBUTIONS

W.L. and H.-o.I. designed the project. W.L. performed all biochemical experiments. Y.T. and H.-o.I. supervised the project. W.L. K.S., M.N., Y.T. and H.-o.I. analyzed data. W.L., Y.T. and H.-o.I. wrote the manuscript with edits provided by all authors. All authors discussed the results and approved the manuscript.

## ACKNOWLEDGEMENT

We thank Atsushi Takeda for providing pCRHA-AGO4 and pCRHA-AGO6. We also thank all the members of the Tomari laboratory for discussion and critical comments on the manuscript, Kaori Kiyokawa for experimental assistance, and Life Science Editors for editorial assistance. This work was supported in part by JST, PRESTO, Japan (grant JPMJPR18K2 to H.-o.I) and JST, FOREST, Japan (grant JPMJFR204O to H.-o.I).

