## Supplementary information for "The mechanisms of siRNA selection by plant Argonaute proteins triggering DNA methylation"

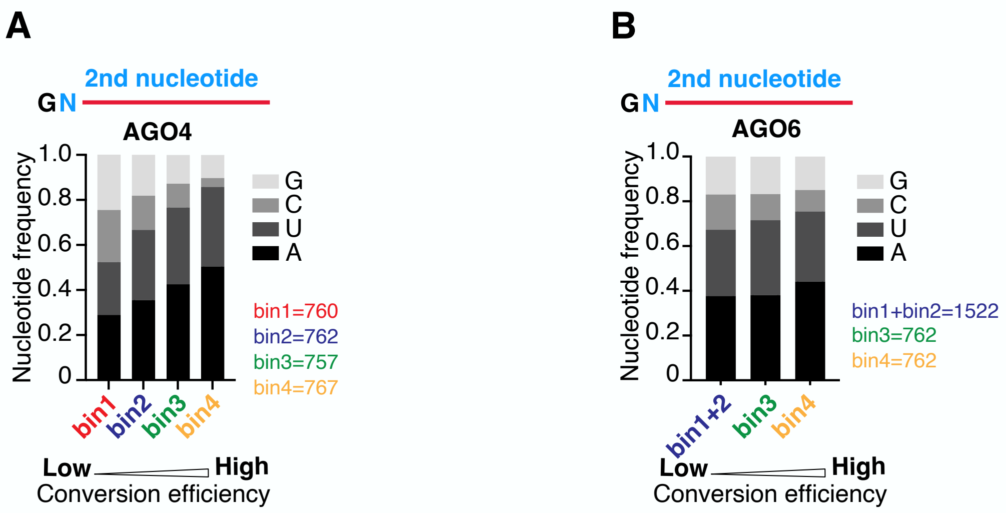


**Supplementary Figure 1. Second nucleotide frequency of AGO4- and AGO6-bound 5′ G small RNAs.**

Second nucleotide frequency of AGO4- (A) and AGO6- (B) bound 5′ G small RNAs. Number of P4RNAs in each bin is shown on the right side. Since those P4RNAs with 0 match to the AGO6-bound 5′ G small RNAs exceed the capacity of a single bin, we merged the bin 1 and bin 2 as one bin. P4RNAs with higher CEs tend to have an A/U nucleotide compared to a C/G nucleotide at the g2 position in both AGO4 and AGO6.

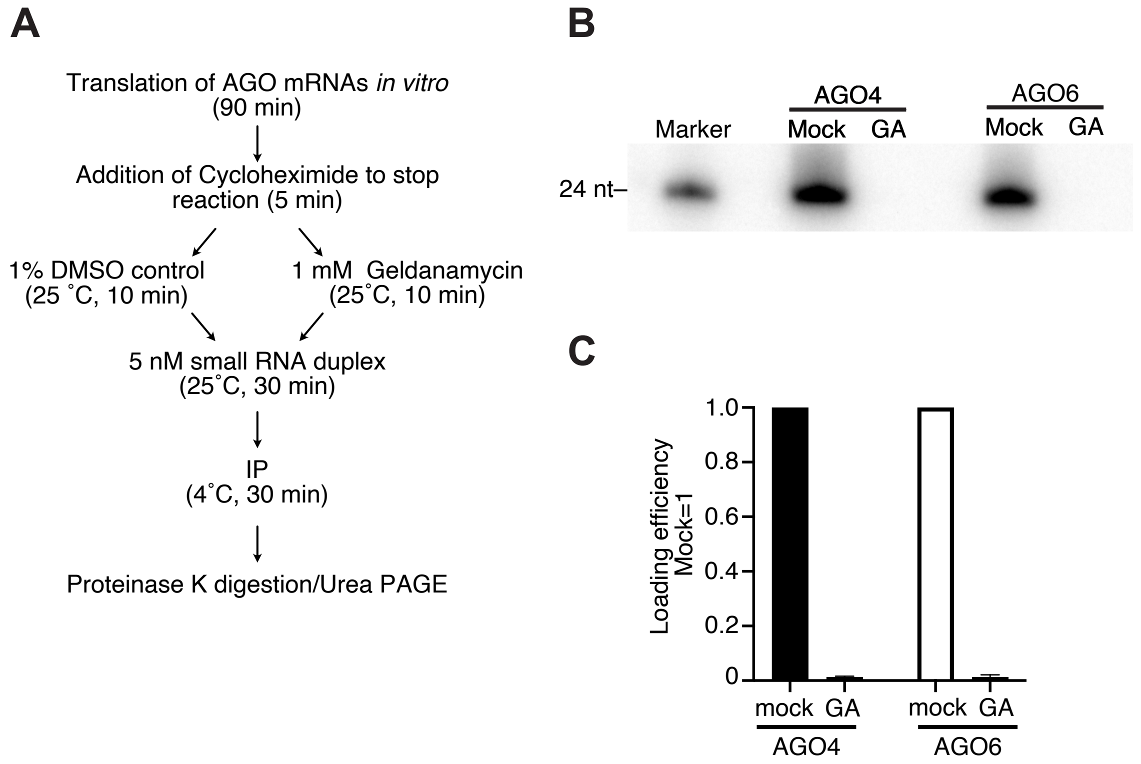


**Supplementary Figure 2. Chaperon inhibitor abrogated loading of small RNAs by AGO4 and AGO6.**

(A) Schematic for the RISC assembly with the chaperone inhibitor geldanamycin or DMSO control.

(B) RISC assembly in BY-2 lysate. HSP90 chaperone inhibitor geldanamycin abrogated loading of small RNAs in both AGO4 and AGO6, comparing to the mock group (DMSO control).

(C) Quantification of loaded siRNAs in (B). The band intensity of siRNAs was normalized to the value of mock group. The graphs show the mean ± SD from three technically independent experiments.

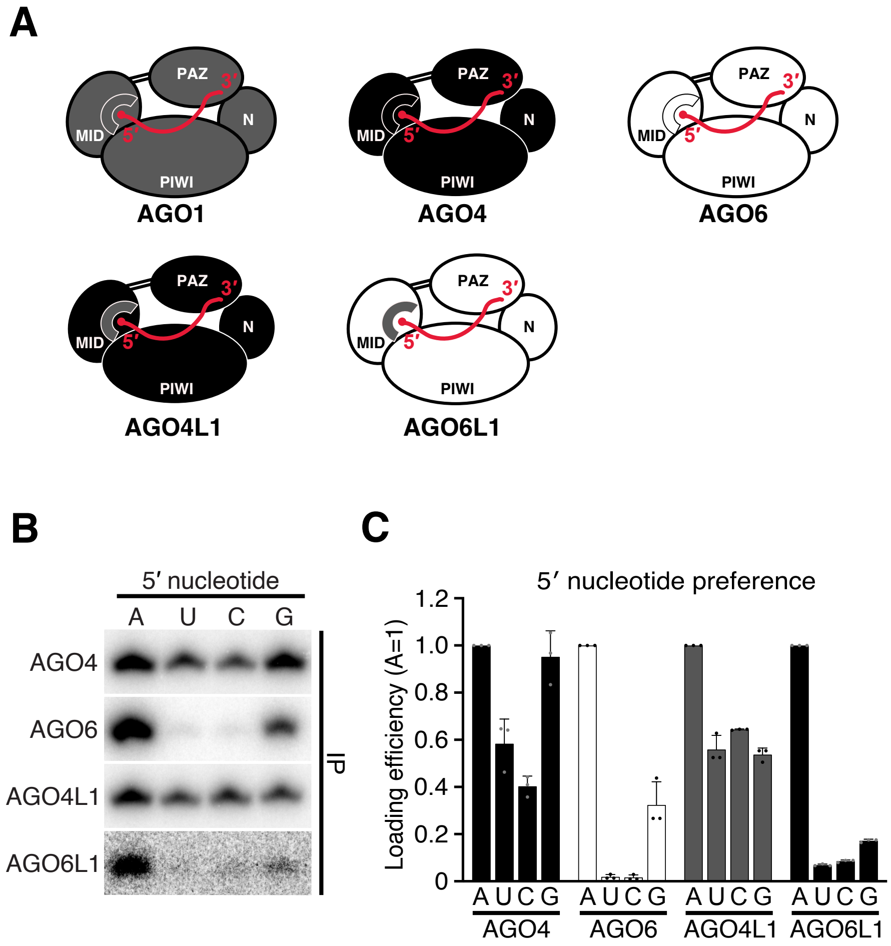


**Supplementary Figure 3. The nucleotide specificity loop in the MID domain does not determine the 5′-nucleotide preference of AGO4 and AGO6.**

(A) Schematic of chimeric AGO4 and AGO6 proteins with AGO1's nucleotide specificity loop.

(B) *In vitro* RISC assembly with chimeric AGO4/6 proteins. AGO4L1 and AGO6L1 showed similar 5′-nucleotide preference as AGO4 and AGO6, respectively.

(C) Quantification of loaded siRNAs in (B). The band intensity of siRNAs was normalized to the value of 5′ A. The graphs show the mean ± SD from three technically independent experiments.
